# Acute threats modulate hunger circuit dynamics to instruct avoidance behaviour

**DOI:** 10.64898/2026.02.17.706314

**Authors:** Alex Reichenbach, Daria J Paulis, Vasilios Drakopoulos, Abigail Wust, Harry Dempsey, Claire J Foldi, Romana Stark, Zane B Andrews

## Abstract

During foraging animals must balance food-seeking with predator avoidance, yet how the brain integrates sensory information relating to food and threat remains unclear. Using in vivo calcium imaging in mice, we show that hunger-sensitive AgRP neurons in the hypothalamus are rapidly inhibited by threats across a threat imminence continuum, from a low environmental risk to a high physical restraint threat, independent of fasting state. This suppression is driven by GABAergic inputs from dorsomedial hypothalamus (DMH) neurons, which increase activity during threat exposure. While AgRP population activity shows uniform inhibition, pathway-specific monitoring using axonal GCaMP reveals distinct projection patterns. For example, AgRP terminals in BNST and LH decrease activity to both threat and food, while threats increased AgRP axonal activity in the PVN. Furthermore, location-specific optogenetic inhibition of AgRP neurons conditions spatial avoidance, mimicking a threat-induced defensive behavioural responses. These findings reveal a hypothalamic circuit where DMH GABA neurons suppress AgRP activity in response to external threats, prioritising avoidance over food-seeking to optimise adaptive behavioural responses during foraging.

## Introduction

Foraging has frequently been used as an experimental model to explore the neural mechanisms underpinning decision-making, as it assesses an animal’s ability to efficiently seek and exploit resources within an environment ^1^. A key factor for optimal foraging is the ability to integrate both interoceptive need states with external sensory information. External sensory information may represent cues predicting resource quality, location, and quantity or may indicate obstacles and dangers associated with obtaining the necessary resources. In the case of foraging, food or water are often the essential sought-after resources, which are driven by the interoceptive states of hunger and thirst ^2^, respectively. Thus, foraging animals may encounter behavioural conflicts driven by competing needs, such as a hungry animal’s need for food as well as the need to avoid external danger or predation. These competing needs create a useful animal model to explore the neural mechanisms underlying approach-avoidance decision making.

This raises some important considerations for the neural control of decision-making during foraging. First, homeostatic circuits of need, using hunger as an example, should affect the salience of external sensory information. Indeed, hunger is a strong survival pressure encouraging motivated food seeking, risk taking, pain tolerance, memory acquisition and exploration in open spaces or unfamiliar terrain to find food ^3–9^. Hypothalamic Agouti-related peptide neurons (AgRP) are key hunger-sensing neurons in the brain ^10^ and states of hunger increase AgRP neural activity and gene transcription ^11,12^. Moreover, AgRP neurons mediate the effects of hunger on behaviour as either chemogenetic or optogenetic manipulation of AgRP neurons drives food intake, motivated food-seeking, exploration, behavioural resilience to stressors, learning and memory function ^7–9,13–18^. Thus, AgRP neurons, as interoceptive hunger-sensing neurons, influence foraging by placing greater weight to approach behaviours during competing need states ^4,9^.

The second consideration for the neural control of decision-making during foraging is how external sensory information affects interoceptive hypothalamic hunger circuits. Over the last decade it has become clear that the sensory detection of food suppresses the activity of AgRP neurons ^12,19,20^. Although fasting increases AgRP neural activity, the sensory detection of food or conditioned food cues immediately suppresses AgRP neuronal activity, where the magnitude of suppression is greatest to palatable calorie dense foods or in hungry animals ^12,19,20^. The sustained suppression of AgRP neural activity signifies sufficient calorie intake and requires gastrointestinal feedback via neuroendocrine feedback from gut peptides or the gut-brain neuroaxis ^21–23^. In the absence of calorie intake, or insufficient calorie intake, AgRP neurons quickly return to higher levels to drive further food seeking behaviour. AgRP neurons encode learned information associated with cues or contexts that predict calorie availability. For example, the presentation of a flavoured gel with calories produces a greater fall in AgRP neuronal activity after the initial first exposure ^23^. Moreover, sustained AgRP neuronal photoinhibition of fasted mice within a context increases food intake when subsequently exposed to the same context at a later date ^7^. In addition to external food cues, different sensory information affects AgRP neural activity including olfactory cues linked to social contact ^24^, circadian cues ^25,26^, cold exposure ^27^, thermal pain3. These studies highlight AgRP neurons provide an internal drive to approach food and external sensory information regulates their activity. Moreover, we recently demonstrated that deletion of ghrelin receptors from AgRP neurons attenuates the sensory response to food cues ^28^. Thus, the appropriate internal sensing of energy state is required for AgRP neurons to respond appropriately to external sensory cues.

When animals forage, food is not the only important external factor influencing behaviour, there is an elevated threat to predation. Indeed, hunger promotes exploration, often into unfamiliar environments, which can be dangerous as food seeking animals are exposed to greater risk of predation. This scenario presents an important approach-avoidance behavioural conflict, where the decision to approach food or avoid a threat source becomes more salient and critical to survival. Interestingly, predator threat using a live rat stress assay rapidly suppressed AgRP neuronal activity while shifting behaviour away from food seeking towards self-preservation ^29^. This suppression of AgRP neuron activity to threat was mediated by a polysynaptic pathway involving CCK neurons in the dorsal premamillary nucleus. However, the identity of the presynaptic GABAergic input onto AgRP neurons remains unknown. Moreover, increased activity of specific AgRP neuron pathways drive food intake, such as AgRP-PVN, PVT, LH, BNST, MeA, MPOA ^14,30–32^, whereas others do not AGRP-PBN, PAG, CeA ^31^. Under predatory threat only the AgRP-BNST and LH, but not AgRP-PVN and PVT promote food seeking ^29^, suggesting subpopulations of AgRP neurons integrate information from different external sensory cues.

The behavioural response to threats exists on a predatory threat imminence continuum ^33^, in which animals select an appropriate defensive behaviour depending on threat proximity. This threat imminence framework starts with low potential threat (pre-threat encounter), such as entering a dangerous zone, where risk assessment and avoidance behaviours are important. Detection of a predator (post-threat encounter) within an environment promotes escape or freeze behaviours depending on context, whereas predator contact during a close encounter (circa-strike) causes defensive attack or tonic immobility to feign death. Here we considered a series of experiments designed to probe AgRP neuronal responses to potential scenarios within this threat imminence framework encountered during foraging, including entering a dangerous zone, detection of a predator and physical capture and restraint. We explored the presynaptic GABAergic inhibitory inputs and explored if AgRP neuron subpopulations, based on pathway projections, respond to different external stimuli. Finally, we used location-specific optogenetic approaches to examine the function of the threat-induced suppression of AgRP neurons.

## Results

### AgRP neuronal response to threat within the threat imminence framework

As mice are a prey species, they naturally avoid open spaces to reduce potential threat ^33^. To test how a low threat affects AgRP neuronal activity, we used a zero maze with defined safe zones (closed arms) and danger zones (open arms) to simulate environmental danger linked a pre-threat encounter during foraging. We prepared *Agrp*-ires-cre mice for *in vivo* fiber photometry with GCaMP7s (Fig 1A) and recorded AgRP neuronal activity as mice moved around a zero maze (Fig 1B). While there were no differences in the number of exit or entry transitions in the closed (safe) zone (Fig 1C), z-score analysis showed an increase and decrease in AgRP neuronal activity compared to baseline when time locked entering or exiting the safe zone respectively (Fig1D,E). Importantly, the change in AgRP neuronal activity was not different in fed or fasted mice when mice entered or exited the safe zone (Fig 1F,G), showing the internal hunger state does not affect how the external environment influenced AgRP activity. To increase the threat potential linked with foraging, we simulated post-encounter threat response using an aerial predator approach towards mice in their home cage (Fig 1H). Aerial predator threat reduced AgRP neuron activity compared to baseline when immediately over the home cage and, similar to the zero maze, the suppression of AgRP neuronal activity did not depend on the internal hunger state (Fig 1I). To simulate a proximal circa-strike threat associated with foraging, we monitored AgRP neuron activity in fed or fasted mice during 15 minutes of restraint stress followed by a 30-minute release period and a 30-minute feeding period. Restraint stress immediately reduced AgRP neuronal activity in both fed and fasted mice compared to baseline and there was no difference in suppression between fed or fasted restraint stressed mice (Fig 1J, Suppl Fig1). Although there was no difference in AgRP neuron activity during the release period compared to the pre-stress baseline (Fig 1J), AgRP neuronal activity increased significantly faster in fasted mice compared to fed mice relative to the 5 mins prior to release (Fig1L). In response to chow consumption, AgRP neuronal activity decreased in both fed and fasted, stressed versus non-stressed mice compared to a pre-stress baseline (Fig 1J). Food intake was significantly increased in fasted compared to fed mice, however, stress had no effect on subsequent feeding 30 minutes after release from restraint (Fig 1K; two-way ANOVA, main effect of metabolic state). When AgRP neural activity during the chow consumption period was compared relative to a 5-minute baseline period prior to chow exposure, there was a significantly greater decrease in activity in fasted mice compared to fed mice, with no differences caused by stress (Fig 1M).

**Figure 1.**
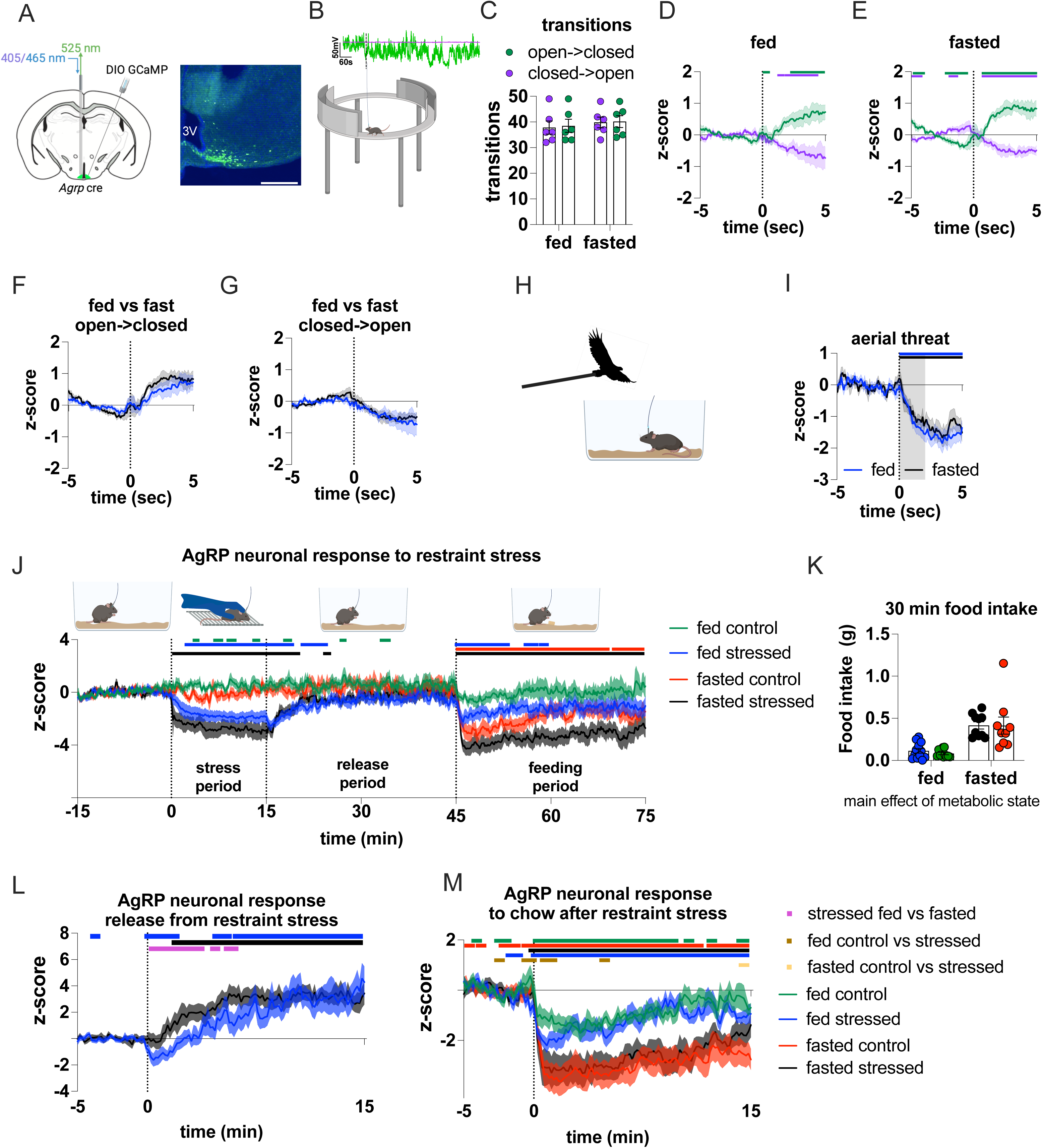
AgRP neurons respond to acute threats A) schematic of viral approach and fiber placement to record neuronal activity of AgRP neurons and representative immunohistological image, scale bar: 500µm. B) Raw trace and experimental setup for fiber photometry recordings during the elevated Zero-maze test. C) Number of transitions from open to closed zones (safety seeking; green dots) and from the closed zones into the open zones (danger signalling; purple dots) of the Zero-maze in fed state and after over-night fast. Average AgRP neuronal activity during transitions from open to closed zones (safety seeking; green) and closed into the open zones (danger signalling; purple) in fed (D) and fasted (E) state show activity changes in opposite direction with no differences for fed (F) or fasted (G) state; N=6. H) Schematic of aerial threat experiment and (I) average AgRP neuronal response to the aerial threat in fed (blue, N=10) and fasted (black, N=6) state and the grey box represents when the aerial threat was above the cage. J) AgRP neuronal activity over 90-minute recording showing the response to 15-minute restraint stress and 30-minute food access relative to the pre-stress baseline period. K) food intake after restraint stress. L) AgRP neuronal activity increase after release and M) in response to chow access relative to the 5-minute period prior to release or chow food presentation respectively; N= 11 fed control; N= 9 fasted control; N= 12 fed stressed; N= 10 fasted stressed. Significant differences are indicated by the lines above graphs as calculated by bootstrapped confidence intervals with 1000 resamples at 95% confidence and 0.5 sec consecutive threshold; changes from baseline are (same coloured) or between groups as indicated. Scale bar: 500µm. For detailed statistical information see Supplementary Table 1.

### DMH GABA neurons provide inhibitory input to AgRP neurons

Next, we examined the source of the presynaptic inhibitory input onto AgRP neurons. As the DMH is a stress responsive nucleus ^34^ and a population of DMH GABAergic neurons provide inhibitory input onto AgRP neurons ^35,36^, we hypothesised that DMH GABAergic neurons inhibit AgRP neuronal activity during threats. To test this hypothesis, we prepared *Vgat* cre mice for photometry to monitor the activity of DMH*^Vgat^* neurons during zero maze, aerial predator approach and restraint stress to model pre and post encounters and the circa-strike period of the threat imminence continuum (Fig 2A-D). DMH*^Vgat^* neuronal activity decreased when entering the closed zone and increased when entering open space in the zero maze (Fig 2B), and in response to an aerial threat (Fig 2C) or restraint stress (Fig 2D) in a manner that opposes AgRP neuron activity to the same threats (Fig 1). There was also a significant increase in DMH*^Vgat^* neuronal activity at the beginning of food consumption consistent with previous reports ^36^. We confirmed that threat responsive DMH*^Vgat^*neurons project to the arcuate nucleus by monitoring DMH*^Vgat^* axonal terminals in the ARC (Fig 2E). Similar to the soma activity of DMH*^Vgat^* neurons, axonal activity in the ARC increased in response to entering an open space and decreased when entering the closed zone in the zero maze (Fig 2F). In addition, axonal activity increased to an aerial threat (Fig 2G) or during restraint stress (Fig 2H). To further ascertain that AgRP neurons receive GABAergic input in response to threat within the threat imminence continuum, we injected *Agrp* cre mice with cre-dependent GABASnFR (Fig 2I). We found that GABA release onto AgRP neurons increased when mice entered the open space of the zero maze (Fig 2J), however no decrease in GABA release onto AgRP neurons was observed when mice entered the closed zone, which contrasts with results from DMH*^Vgat^* neuron soma activity or axonal activity within the ARC. Both aerial predator approach and restraint stress increased GABA release onto AgRP neurons (Fig 2K, L), consistent with DMH*^Vgat^* neuron axonal activity within the ARC and temporally aligning with the suppression of AgRP neurons in response to the threats (Fig 1). While these studies show increases in DMH*^Vgat^* neuronal activity and GABA release correlate with AgRP suppression, they do not show causality. To this end, we first used flp-dependent GABASnFR with the cre-dependent inhibitory opsin JAWS in *Vgat* cre::*Agrp* flp mice (Sup Fig 2A) and showed that inhibition of DMH*^Vgat^* neurons directly reduced GABAergic input on AgRP neurons at baseline and during restraint stress (Sup Fig 2B,C). To demonstrate that DMH*^Vgat^*neurons are functionally necessary to inhibit AgRP neurons in response to threats within the threat imminence framework, we injected *Vgat* cre::*Agrp* flp mice with the cre-dependent inhibitory opsin JAWS in the DMH and flp-dependent GCaMP6f targeting AgRP neurons (Fig 2M). In response to an aerial predator threat without inhibiting DMH*^Vgat^* neurons, AgRP activity decreased as expected (Fig 2N), However, DMH*^Vgat^* neuronal inhibition during an aerial threat surprisingly did not just prevent the expected decrease in AgRP activity, it resulted in an increase in AgRP activity (Fig 2N). These results suggest removing the inhibitory input from DMH*^Vgat^* neurons during aerial threat unmasks and allows an excitatory glutamatergic input to increase AgRP activity unopposed. To show that DMH*^Vgat^* neurons cause AgRP neuron inhibition during restraint stress, we first restrained mice and observed a decrease in AgRP neural activity. However, during restraint, inhibition of DMH neurons with JAWS returned the activity of AgRP neurons to baseline (Fig 2O). These optogenetic inhibition studies demonstrate that DMH*^Vgat^* neurons increase GABA binding to AgRP neurons and inhibit AgRP neurons at baseline and during an aerial threat and restraint stress.

**Figure 2.**
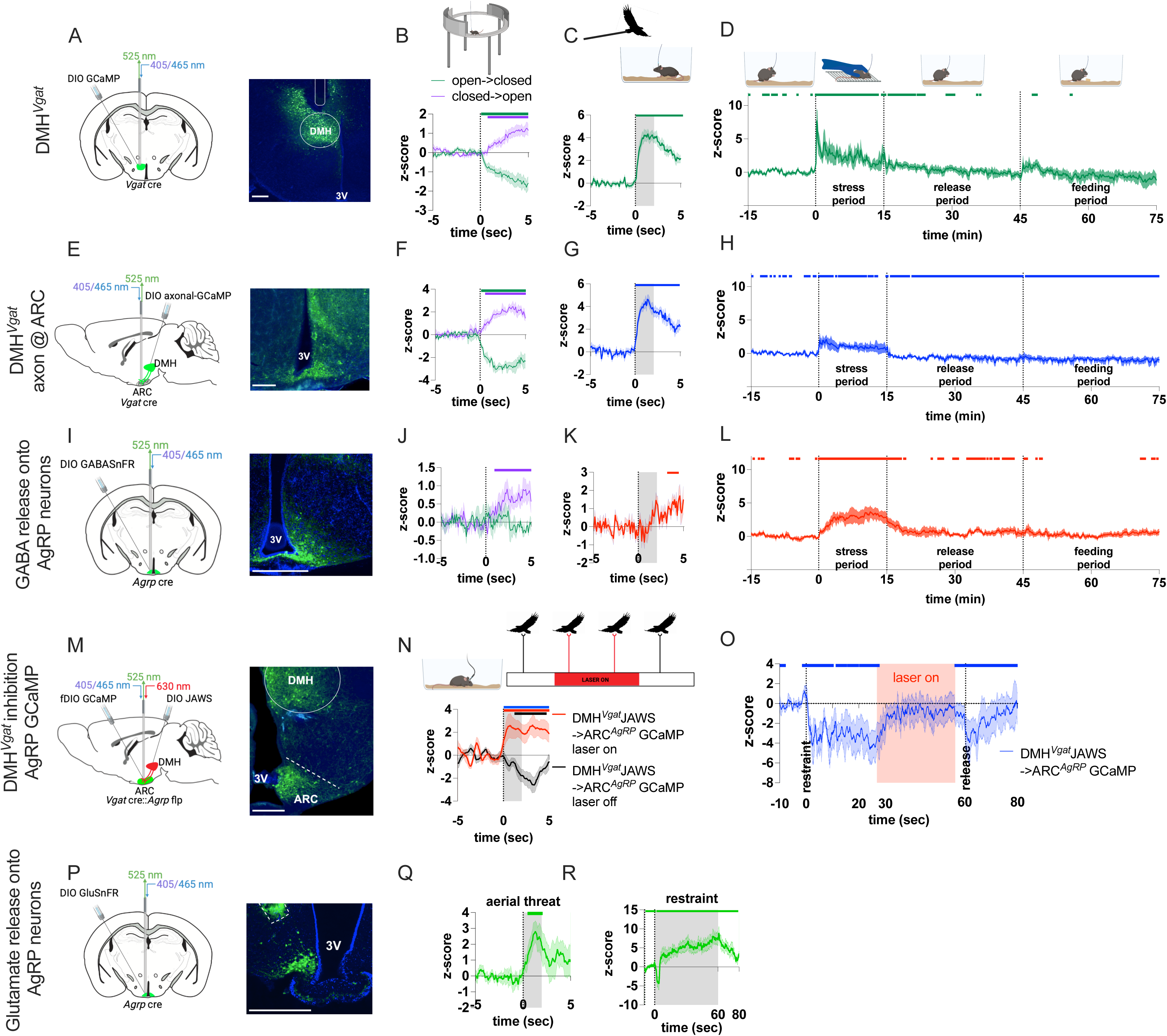
DMH^Vgat^ neurons input AgRP neurons during acute threats. Schematic of viral approach and representative image used to monitor DMH^Vgat^ neurons (A). Average change in activity of DMH^Vgat^ neurons (B) during the transition from open to closed zones (safety seeking; green) and from closed to open zones (danger signalling; purple) in response to aerial threat (C) and in response to 15-minute restraint stress and 30-minute chow access (D). Schematic of viral approach and representative images used to monitor DMH^Vgat^ axonal projections within the ARC (E). Average change in DMH^Vgat^ axonal activity within the ARC during the transition from open to closed zones (green) and from closed to open zones (purple) (F), in response to aerial threat (G) and in response to 15-minute restraint stress followed by 30-minute chow access (H). Schematic of viral approach and representative images used measure GABA release onto AgRP neurons (I). Average change in GABA binding to AgRP neurons during the transition from open to closed zones (green) and from closed to open zones (purple) (J), in response to aerial threat (K) and in response to 15-minute restraint stress and 30-minute chow access (L). Schematic of the viral approach to inhibit DMH*^Vgat^* neurons while recording AgRP neuronal activity (M). AgRP neuronal response to aerial threat with and without optogenetic DMH*^Vgat^* neuron inhibition (N) and AgRP neuronal response to one minute restraint with 30 second DMH*^Vgat^* neuronal inhibition starting 25 seconds after restraint onset (O), as indicated by the pink box. Schematic of viral approach to measure glutamate release onto AgRP neurons (P) and glutamate release in response to aerial threat (Q) and one minute restraint (R). N(DMH*^Vgat^*) = 7; N(DMH*^Vgat^* axons @ARC) = 3, N(GABA@AgRP) = 3, N(DMH*^Vgat^*JAWS) = 5, N(Glutamate@AgRP) = 3. Significant differences are indicated by the lines above graphs as calculated by bootstrapped confidence intervals with 1000 resamples at 95% confidence and 0.5 sec consecutive threshold; changes from baseline are indicated with the same coloured. Grey boxes in C,G, K, N and Q indicate when the aerial threat was above the cage or the restraint period (R). Scale bar: 500µm. For detailed statistical information see Supplementary Table 1.

To test the hypothesis that removing the inhibitory input from DMH*^Vgat^* neurons uncovers an excitatory glutamatergic input, we employed a cre-dependent glutamate sensor in *Agrp* cre mice (Fig 2P). We detected an acute increase in glutamate release onto AgRP neurons to an aerial threat (Fig 2Q) and a prolonged increase during restraint, which persisted until mice were released after 60 seconds (Fig 2R). To examine the neural source of glutamate release onto AgRP neurons, we injected *Vglut2* cre::*Agrp* flp mice with cre-dependent axonal GCaMP in PVN*^Vglut^*^2^ or LH*^Vglut^*^2^ neurons and a flp-dependent red-shifted calcium indicator RGECO to target AgRP neurons (Sup Fig 2D). This approach enabled to us simultaneously monitor PVN*^Vglut^*^2^ or LH*^Vglut^*^2^ axonal activity within the ARC and AgRP neuronal activity. PVN*^Vglut^*^2^ axonal activity within the ARC increased to both aerial threat and restraint stress (Sup Fig2 E-F), however, LH*^Vglut^*^2^ axonal activity within the ARC decreased to both threats. These results suggest that the source of glutamatergic input onto AgRP neurons stems from PVN*^Vglut^*^2^ neurons, where a direct monosynaptic input from PVN*^Vglut^*^2^ neurons to AgRP neurons has been described previously ^37^.

### Specific AgRP neuronal projection pathways respond to different sensory information

It is possible that different AgRP neuronal subpopulations respond to different glutamatergic and GABAergic presynaptic input in response to different sensory information. Alternatively, DMH*^Vgat^* neuronal inhibition could differentially affect the activity of AgRP neuronal subpopulations. To address this possibility and circumvent the drawback of homogenous sampling of AgRP neuronal population activity, we used axonal GCaMP to monitor AgRP axonal activity at different projection sites. In response to food presentation in fasted mice, AgRP neuronal activity was significantly decreased compared to baseline in all terminal regions examined, including the PVN, LH, BNST, PVT and PAG (Fig 3 D, H, L, P, T). We then probed the responses to an aerial threat and restraint stress, as within the threat imminence framework aerial threat (post-threat encounter) and restraint stress (circa-strike) often elicit different defensive behaviours, such as freezing or escape (aerial threat) as compared to defensive attack or tonic immobility (restraint stress) ^38,39^. AgRP axonal activity within the BNST and LH significantly decreased compared to baseline in response to aerial threat or restraint stress (Fig 3E-F, I-J). However, we observed an increase in AgRP axonal activity in the PVN to both aerial and restraint stress (Fig 3M, N), despite a decrease to food presentation. To confirm this, we injected a cre-dependent retrograde AAV GCaMP in the PVN and a redshifted cre dependent RGECO into the arcuate nucleus of *Agrp*-ires-cre mice (Supp Fig 3A). This allowed simultaneous recording of overall AgRP neuronal response and the subset of AgRP neurons projecting to the PVN. Similar to axonal activity in the PVN, the AgRP-PVN pathway was activated by aerial threat and restraint stress (Supp Fig 3B,C), even though the same stimuli decreased the AgRP population response (Supp Fig 3B,C). This suggests the PVN receives different AgRP neural input in response to food as compared to threat, which is likely a reflection of different presynaptic input onto AgRP neurons. In addition, AgRP axonal activity in the PVT and PAG showed differential responses to the different threats. In the PVT, aerial threat caused a rapid and transient decrease in activity while the aerial threat is close, followed by an immediate increase post threat period (Fig 3Q). Restraint stress resulted in an initial decrease followed by ramping in activity that returned to baseline after release (Fig 3R). In the PAG, aerial threat increased, whereas restraint stress decreased AgRP axonal activity (Fig 3U,V). These results collectively show that AgRP neurons do not respond homogenously to threats as it is described for food presentation and AgRP neuron projections respond differently to aerial threat or restraint stress, suggesting a graded response depending on the imminence of the threat.

**Figure 3.**
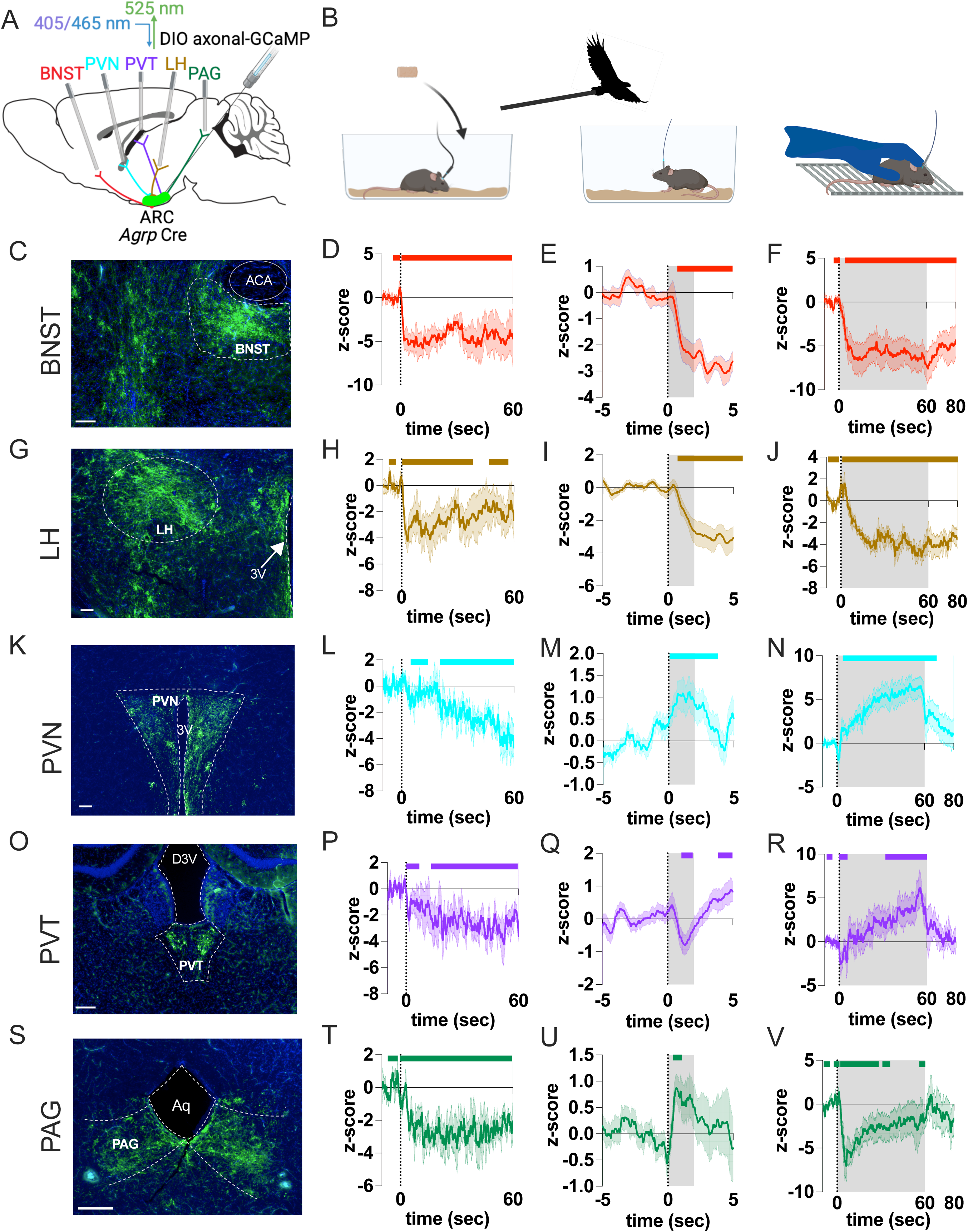
AgRP axonal projections show differential response to acute threats. A) Schematic of viral approach to target AgRP axonal projections using axonal GCaMP in the bed nucleus of the stria terminals (BNST), lateral hypothalamus (LH), paraventricular nucleus of the hypothalamus (PVN), paraventricular nucleus of the thalamus (PVT), and periaqueductal gray (PAG). Each animal had no more than two implant sites. B) Schematic images to represent the experiments conducted; food in cage, aerial threat and restraint stress. Axonal GCaMP expression in the BNST (C) and AgRP axonal activity in the BNST in response to food presentation (D), aerial threat (E) and acute restraint stress (F). Axonal GCaMP expression in the LH (G) and AgRP axonal activity in the LH in response to food presentation (H), aerial threat (I) and acute restraint stress (J). Axonal GCaMP expression in the PVN (K) and AgRP axonal activity in the PVN in response to food presentation (L), aerial threat (M) and acute restraint stress (N). Axonal GCaMP expression in the PVT (O) and AgRP axonal activity in the PVT in response to food presentation (P), aerial threat (Q) and acute restraint stress (R). Axonal GCaMP expression in the PAG (S) and AgRP axonal activity in the PAG in response to food presentation (T), aerial threat (U) and acute restraint stress (V). N=4 per axonal projection site. Grey boxes indicate when the aerial threat was above the cage (E,I,M,Q,U) or the restraint period (F,J,N,R,V). Significant differences are indicated by the lines above graphs as calculated by bootstrapped confidence intervals with 1000 resamples at 95% confidence and 0.5sec consecutive threshold; changes from baseline are indicated with the same coloured. Abbreviations: ACA, anterior commissure; Aq, cerebral aqueduct; D3V, dorsal third ventricle; 3V, third ventricle. Scale bar: 200µm. For detailed statistical information see Supplementary Table 1.

### AgRP neuronal inhibition promotes conditioned avoidance

Our results show that all threats examined within the threat imminence framework suppress AgRP population responses. To assess the physiological function of this threat-induced inhibition, we designed an experiment to specifically test whether the fall in activity is sufficient to promote conditioned avoidance. We injected a red-shifted cre-dependent inhibitory opsin (JAWS) in the arcuate nucleus of *Agrp*-ires-cre and WT litter mice (Fig 4A). We placed mice into a Y-maze and photoinhibited AgRP neurons every time mice entered the left arm (Fig 4B). This was referred to as the paired arm, as AgRP neuron photoinhibition was paired with left arm entry. We trained mice for 5 days to associate the paired arm of a Y-maze with AgRP neuronal inhibition (Fig 4B). On the test day, mice were placed in the Y-maze without optogenetic modulation and behaviour recorded. Although the time to first entry in the paired or control arm was not different between *Agrp* cre and WT mice (Fig 4C), *Agrp* cre mice made significantly more entries into the control arm compared to WT mice (Fig 4D). Importantly, *Agrp* cre mice spent significantly less time in the arm paired with photoinhibition compared to the control arm, indicating AgRP neural inhibition promoted conditioned avoidance of the paired arm. Consistent with this idea, *Agrp* cre mice spent less time per bout, moved faster with less stationary time in the arm paired with AgRP neuron photoinhibition compared to the control arm (Fig 4F-H). Together, this suggests that *Agrp* cre mice learned to associate the photoinhibited arm with a sense of threat that produces vigilant and avoidant behaviour in this area.

**Figure 4.**
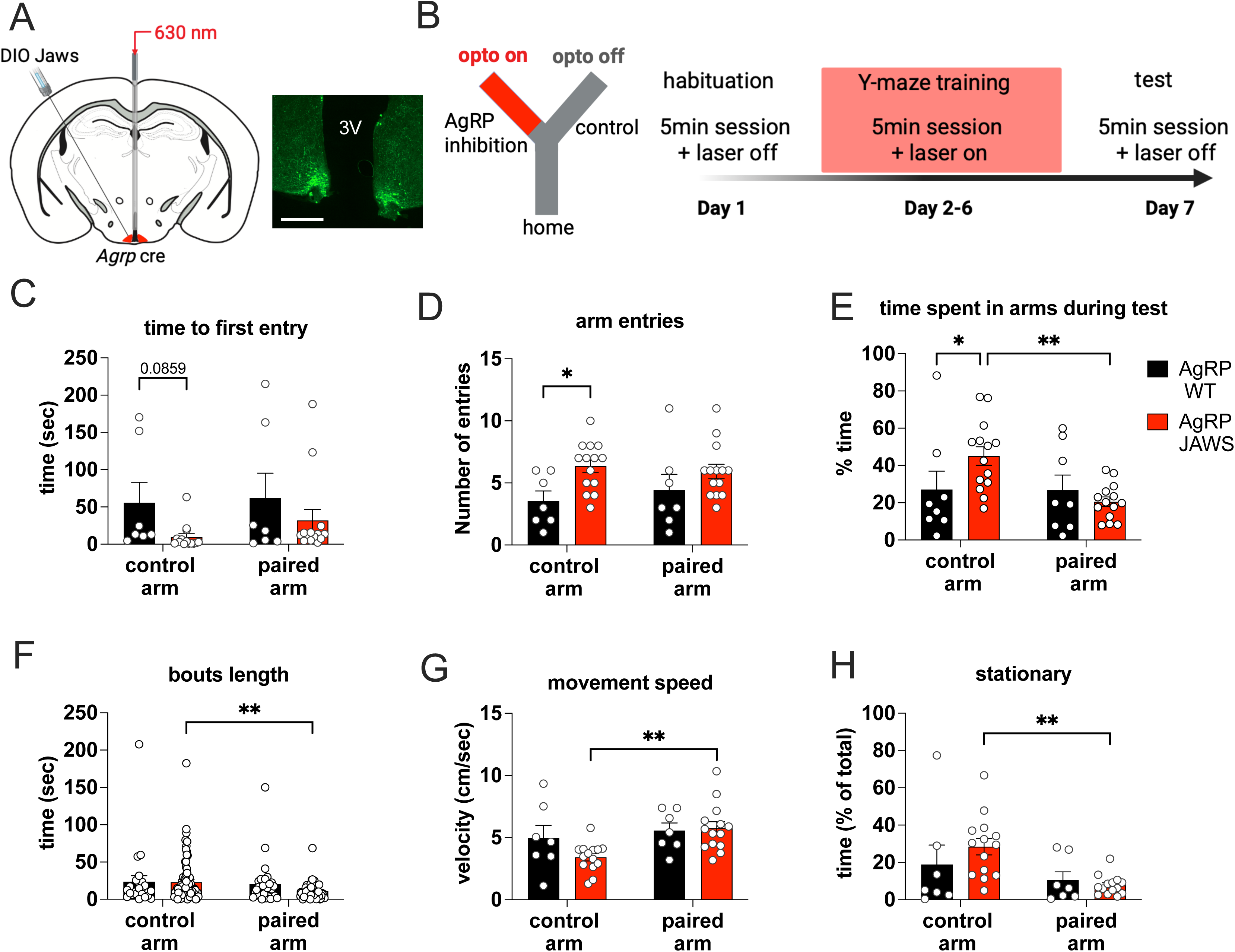
AgRP photoinhibition induces context-specific conditioned avoidance. A) Schematic of viral approach using cre-dependent JAWS to inhibit AgRP neurons in *Agrp* cre and WT mice; scale bar = 200µm. B) Experimental set up and protocol used to train mice to associate one arm of a Y-maze with AgRP photoinhibition. During training days, behavioural recognition software triggered AgRP photoinhibition when mice entered one arm of the Y-maze (paired arm) but not the other arm (control arm). No photoinhibition occurred during the test. Although there was no significant difference in time to first entry into control arm and paired arm (C), AgRP JAWS had a significantly increased the number of control arm entries compared to AgRP WT mice (D). AgRP JAWS mice also spent significantly less time (E) with a shorter bout length (F) per entry in the arm paired with AgRP photoinhibition compared to AgRP WT mice. AgRP JAWS mice had increased movement speed (G) and reduced stationary time in the paired arm (H). N= AgRP WT 7 AgRP JAWS = 14. Data expressed as mean +/- SEM and statistical analysis was performed using 2-way ANOVA, * indicates p<0.05, ** p<0.01. For detailed statistical information see Supplementary Table 1.

## Discussion

Foraging requires the ability to respond to interoceptive signals of hunger and sensory information within an external environment. Thus, hunger-sensing AgRP neurons are ideally positioned to optimise foraging, as they respond to internal signals of hunger, such as ghrelin 28, and respond to the sensory detection of food and conditioned food cues ^20,35^. However, when foraging, animals need to balance effective food seeking while avoiding external threats and danger. In this study, we examined how AgRP neurons respond to potential foraging threats within a threat imminence continuum ^33^, starting with a low pre-encounter threat (zero maze), a post-encounter threat (aerial threat) and a high circa-strike threat (restraint stress). We found that all threats acutely inhibited overall AgRP neuron population activity regardless of threat potential within the threat imminence framework. Unlike AgRP neuronal responses to food or food-predicting cues ^12,19,20^, fasting did not affect the sensitivity to threat detection along the threat imminence continuum, which is not surprising since the potential external threat of predation does not depend on the internal hunger state of the potential prey. We demonstrated that DMH*^Vgat^* neurons inhibit AgRP neuronal activity and increase GABA release onto AgRP neurons during threat exposure. Moreover, monitoring AgRP axonal activity in the LH, BNST, PVN, PVT, PAG demonstrated a decrease to food presentation in all regions examined. However, only axonal activity in BNST and LH decreased to both aerial threat and restraint stress. AgRP axonal activity in the PVN increased to both threats, whereas the PVT and PAG showed different responses to aerial threat or restraint. Thus, not all AgRP neuronal subpopulations, based on their axonal projections, respond the same to external threats suggesting differences in presynaptic input. Finally, AgRP neuronal photoinhibition in a discrete arm of the Y-maze conditioned subsequent avoidance of that arm, suggesting suppression of AgRP neurons to threat helps mice avoid areas previously associated with danger, thus optimising foraging behaviour.

Our studies highlight that external threats cause acute activation of DMH*^Vgat^* neuronal activity broadly consistent with previous observations using protein markers of DMH activation in respond to threat ^40^. Moreover, photostimulation of DMH inhibitory neurons induces a negative arousal state associated with defensive behaviours ^34^, consistent with the conditioned avoidance observed after AgRP neuronal photoinhibition in our Y-maze test. Microinjections of excitatory amino acids N-methyl-d-aspartic acid (NMDA) in the DMH drive defensive behaviours ^41^, suggesting a multi-synaptic circuit in which glutamatergic neurons activate DMH*^Vgat^* neurons to inhibit AgRP neurons in response to threat. LH*^Vglut^*^2^ neurons are one possibility since they regulate AgRP neurons via a DMH*^LepR^* GABA neuronal input ^35^. Furthermore, LH*^Vglut^*^2^ neurons not only suppress food intake ^42–44^, they also promote avoidance and safety seeking via a variety of specific pathway projections to the tail of the striatum ^6^, lateral habenula ^44^, ventral tegmental area ^42^, the PVN ^45^ and pedunculopontine nucleus ^46^. Moreover, a recent study showed that LH*^orexin^*neurons inhibit AgRP neurons indirectly via a DMH*^Vgat^* inhibitory input, which increases innate avoidance behaviour ^47^. It is important to note that previous studies suggest DMH*^LepR^* GABA neurons directly innervate AgRP whereas DMH*^Vgat^* neurons may also target POMC neurons ^36,48^. Since we used *Vgat* cre mice there is a possibility that some POMC neurons may have received inhibitory input from DMH*^Vgat^* neurons. However, our results do not support this idea since photoinhibition of DMH^Vgat^ neurons prevented the decrease in AgRP activity during restraint stress and directly reduced GABA release on AgRP neurons at baseline and during restraint stress. Moreover, studies showed that restraint stress increases ARC*^POMC^* neuronal firing ^49^ and cfos staining ^50^, suggesting that, at least the majority of POMC neurons in the ARC, are not inhibited by threat-associated input from DMH*^Vgat^* neurons. Nevertheless, POMC neurons and downstream MC4R neurons in the PVN play crucial roles in satiety and body weight so future studies should examine the response of POMC neurons to threat.

It is unusual, and perhaps surprising, that vastly different external sensory information, such as palatable food and restraint stress, both inhibit AgRP neurons and that the source of this inhibition can be traced back to increased GABA inhibitory activity in the DMH ^36^. The similar nature of this input organisation implies that differences in AgRP output pathways may be responsible for divergent behavioural responses to various external stimuli, such as food vs threat. Indeed, activation of different AgRP neuronal projection regions induces a vast array of different behavioural and physiological responses. For example, photostimulation of AgRP terminals in the LH, BNST, PVT, PVN, MeA, MPOA all increase food intake ^14,30–32^, whereas stimulation of AgRP terminals in the dorsal raphe suppresses energy expenditure and thermogenesis ^51^ and in the PBN prevents meal termination ^52^ and inflammatory pain ^3^. In addition to inducing feeding, a AgRP->MeA->BNST, but not AgRP->PVN, pathway suppresses territorial aggression and contextual fear ^32^, while the AgRP->MPOA promotes nest building ^53^ but acutely suppresses parenting behaviour to prioritise food seeking ^30^. In line with different AgRP pathways controlling different behaviours, we identified distinct pathway-specific patterns of AgRP axonal activity in the BNST, LH, PVN, PVT and PAG. Only the LH and BNST showed suppression of activity to food presentation, aerial threat and restraint stress. Intriguingly, photostimulation of AgRP->LH and AgRP->BNST pathways promoted food seeking and intake in the presence of a rat predator ^29^, suggesting these pathways dampening threat avoidance, possibly by inhibiting LH*^Vglut^*^2^ output projections that promote avoidance ^6,42,44,46^. Indeed, our studies support this idea, since inhibiting AgRP neurons in a Y-maze arm during training promoted conditioned avoidance of that same paired arm during testing.

Although food presentation suppressed AgRP axonal activity in the PVN, aerial threat and restraint stress increased axonal activity in the PVN. Both predator and restraint stress activate the HPA system and increase plasma corticosterone ^29,54^ to promote the appropriate behavioural response to the threatening situation. Moreover, previous experiments show that AgRP neuronal activity is required for fasting-induced CORT release, by inhibiting a BNST*^Vgat^* projection in the PVN ^25^. Our results align with this study as aerial threat or restraint stress both increase AgRP axonal activity in the PVN, which would promote HPA activation and CORT release. The fact that food suppresses, whereas threats increase, AgRP->PVN axonal activity suggest different functions of this same pathway in response to different environmental stimuli.

Our studies show AgRP-PVT projections exhibit different responses to threats compared to food. As AgRP neurons are inhibitory, increased activity of AgRP axons in the PVT, which is known to drive food intake ^31^ and promote visual, spatial and contextual cue associations with food availability ^8,55^, would be associated with suppressed PVT activity. Interestingly, rewarding stimuli suppress PVT neurons projecting to the NAc ^56^, whereas arousing and aversive stimuli activate PVT neurons ^57–60^. We suggest that increased AgRP axonal inhibitory activity in the PVT in response to aerial threat and restraint stress may dampen aversive avoidance signaling and facilitate food approach, although future experiments are required to explore this idea.

The PAG is essential to control defensive behaviours ^33^ with a particular role in post-encounter freezing or escape behaviours ^61^, although a recent study suggested an anterior hypothalamic nucleus-PAG pathway is critical for defensive attack during a circa strike phase ^62^. However, a post-encounter aerial threat increased, whereas a circa strike restraint threat decreased AgRP->PAG axonal activity, suggesting AgRP neural activity in the PAG may produce different behavioural outcomes depending on threat imminence and context. Future studies are necessary to examine how the AgRP->PAG pathway controls the expression of defensive behaviours.

Thus, despite aerial threat and restraint stress suppressing average AgRP population activity, we observed distinct differences in AgRP axonal activity that support multifunctional roles for different AgRP neuron pathway-specific subpopulations. One limitation with the use of axonal GCaMP is the potential for local mechanisms to regulate presynaptic input independent from cell bodies. However, injection of retro cre-dependent GCaMP in the PVN of *Agrp*-ires-cre mice produced similar results to axonal GCaMP, further supporting the hypothesis of different AgRP subpopulations responding to different external stimuli. Importantly, simultaneous recordings of AgRP->PVN projecting cell bodies and total AgRP cell bodies show opposing results, suggesting existing AgRP population recordings do not capture sufficient neuronal detail. Future studies with single-cell imaging or pathway specific transcriptomic techniques are required to further elaborate on the existence and roles of AgRP subpopulations.

We propose the suppression of AgRP neuronal activity acts as a salient “stop” signal in response to external events, such as finding food or detecting local threats. In the case of detecting food, this salient “stop” signal allows food consumption, whereas in response to threat and in the absence of food detection, the “stop” signal facilitates vigilance and avoidance. Interestingly, following the suppression of AgRP neuronal activity to food detection, normal feeding behaviour requires sequential activation of LH*^Vgat^* and DR*^Vgat^* neurons ^63^. This observation suggests the coordinated recruitment of downstream neuronal populations may dictate the appropriate behavioural response to the initial sensory-driven inhibition of AgRP neurons, whether it be food or threat. Indeed, our results support this idea since the collective AgRP axonal response (LH, BNST, PVN, PVT, PAG) to threat is distinct from that of food. Moreover, unique AgRP axonal response patterns further distinguish aerial threat from restraint stress suggesting axonal response patterns in different downstream areas may dictate certain behavioural or physiological outcomes.

Furthermore, AgRP axons in the PVN, PVT and PAG respond differently to external sensory information related to threat and food, therefore, we predict that different presynaptic sensory input onto the same AgRP subpopulation may be responsible. While DMH*^Vgat^* neurons provide inhibitory input onto AgRP neurons in response to both food and threat, the relative glutamatergic input may help differentiate food from threat and/or the nature of the threat signal. Indeed, the activity of PVN and LH glutamatergic terminals in the ARC shows distinctive responses during aerial threat and restraint stress. Future studies are required to examine the relative contributions of glutamate and GABA release onto AgRP neurons in response to sensory information.

In summary, we examined how AgRP neurons respond to potential foraging threats within a threat imminence continuum ^33^, starting with a low pre-encounter threat, a post-encounter threat and a high circa-strike threat. We used a functional in-vivo circuit mapping approach and uncovered that AgRP neurons receive GABAergic from DMH*^Vgat^* neurons to help integrate threat signals. We also observed an underappreciated role for glutamatergic input onto AgRP neurons in response to threat detection. Despite threats suppressing AgRP population responses, we surprisingly found various AgRP axonal projections respond differently to a variety of threats. Our studies provide experimental evidence for the existence of AgRP subpopulations and suggest different presynaptic input wiring may underpin the activity of distinct AgRP subpopulations.

## Methods

### Animals

All experiments were conducted in compliance with the Monash University Animal Ethics Committee guidelines (ethics protocol 37004). Male and female mice were kept under standard laboratory conditions with free access to food (chow diet, catalog no. 8720610, Barastoc Stockfeeds, Victoria, Australia) and water at 23C in a 12-hr light/ dark cycle and were group-housed to prevent isolation stress. All mice were aged 8 weeks or older for experiments. *AgRP*-ires-cre(AgRPtm1(cre)Low/J (stock no. 012899), *Vgat* cre::*Agrp* flp and *Vglut2* cre::*Agrp* flp mice were obtained from Jackson Laboratory. Fasted mice had chow removed from food hopper the day before experiments and were refed after completing the experiment.

### Surgery

Viruses for viral infection were purchased from Addgene: AAV9 pGP-AAV-syn-FLEX-jGCaMP7s-WPRE (#104491), AAV1 pAAV.hSyn-FLEX.iGABASnFR.F102G (#112164), AAV1 pAAV.hSyn.FLEX.iGluSnFR3.v857.PDGFR.codonopt (#175180), AAV9 FLEX axonal-GCaMP6 (#112010), AAV8-Ef1a-Coff/Fon-sRGECO (#137127), AAV8-Ef1a- Con/Foff 2.0-sRGECO (#137128), rgAAV-syn-FLEX-jGCaMP8f-WPRE AAV2/9 (#162379), and BrainVTA: rAAV-EF1a-DIO-Jaws-EGFP-ER2-WPRE-hGH polyA (#PT-0103), rAAV- EF1a-fDIO-iGABASnFR-WPRE-hGH polyA,AAV2/9 (#PT-3324). Black fiber optic cannulae for fiber photometry with 1,25mm diameter and 400um core, 0.5NA were purchased from RWD.

Mice were anaesthetized and placed into a heated stereotaxic platform (Stoelting). A approx. 1 cm incision to expose bregma and lamda was cut in AP direction, scull cleaned with 1% hydrogen peroxide and holes drilled over target coordinates. Virus (neat, 200nl for fiber photometry, 100nl for optogenetics) was injected with a pulled glass pipette (at 40nl/min) and pipette slowly withdrawn 5 minutes after infusion was complete and fiber inserted above the injection site targeting AgRP neurons (AP: -1.6; ML:+/-0.2; DV: -5.8) or AgRP axonal projections were targeted with following coordinates from bregma: BNST: AP +0.1 ML -1.3 @10degree DV -4.57; LH: AP -1.2 ML +1.6 @6degree DV -4.9; PVN: AP -0.94 ML +1.2@10degree DV -4.7; PVT: AP -1.3 ML -0.5 @10degree DV -2.7; PAG: AP -4.6 ML -0.75 @10degree DV -2.55. *Vgat* Cre mice received viral injections to target DMH at following coordinates: AP:-1.8; ML:+/- 0.4;DV:-5.2 and fibers were implanted above DMH for GCaMP7s photometry or ARC coordinates for axonal GCaMP6.*Vgat* cre/*AgRP* flp mice used for optogenetic control experiments received 100nl DIO JAWS injection at DMH and 200nl fDIO GCamp or fDIO-iGABASnFR and fiber at ARC coordinates. *Vglut2* cre/*AgRP* flp mice received viral injections to target Vglut2 neurons at the PVN or LH with axonal-GCaMP6 and CreOFFflpON RGECO and fiber targeting the ARC.

All mice had at least 2 weeks recovery from surgery before start of habituation and experiments. Fiber photometry recordings were performed using a TDT fiber photometry rig RZ5 as described before ^17^, or a RWD R821 tricolor multichannel fiber photometry system for axonal GCaMP recordings and combined optogenetic and fiber photometry experiments. For experiments with RWD 821 fiber photometry system 410/470/590nm signal was recorded at 60Hz with 14ms exposure and light set to 5% for 410nm, 19% for 470nm and 23% for 590nm. Fed or over-night fasted mice were tethered to the fiber optic patch cord and acclimatised in their home cage for 5-10 min before recording.

Fiber photometry analysis: Pre-processing of TDT RZ5 raw signals was performed as described before using in house fiber photometry GUI (github/Andrews-Lab/Fiber_photometry_analysis). Briefly, raw signals were down sampled and fitted to 405 isosbestic control and df/f z-scored using pre-stimulus time as baseline. For RWD recordings pre-processing was performed through RWD software (V2.0.0.44140): smoothing coefficient W=15, and exponential baseline fitting coefficient b=8 and df/f z-scored using pre-stimulus time as baseline to match TDT data.

Elevated Zero Maze: At least two minutes baseline in home cage were recorded before fed or fasted mice were lifted onto the open zone of the elevated zero maze and recorded for 10 minutes. Transitions between open and closed zones were timestamped post recording for further analysis.

Looming object: At least one minute baseline was recorded before a looming object (Toothless^TM^ toy dragon, approx. 60cm wingspan) was briefly (approx. 2 seconds) moved over the home cage. This procedure was repeated 3-4 times about 1 minute apart.

Restraint stress: At least 15 minutes baseline were recorded before mice were lifted onto a cage lid and physically restraint (holding behind the head with one hand and the tail with the other hand). After 15 minute restraining, the animal was placed back in the home cage to recover for 30 minutes before one chow pellet was placed into the cage. Food intake was measured 30 minutes thereafter.

Mice undergoing one minute immobilisation were habituated to the fiber photometry tether for 5 min or as long as necessary to stabilise baseline before being restraint in the same fashion as described above inside their home cage.

For optogenetic experiments with inhibitory opsin, mice received constant laser illumination (630nm; 6mW at the fiber tip). For looming object experiment this illumination lasted for 3 minutes, for short immobilisation experiments 30 seconds. For inhibition during rest in home cage, mice received 3 times 1 minute illumination every 3 minutes

YMAZE: We trained *AgRP* Cre / WT control mice infected with cre-dependent JAWS to associate one arm of the maze with AgRP inhibition using closed loop optogenetics with RWD 821 fiber photometry system coupled to 630nm optogenetic laser with ROI based activation based on real-time location of the animal. Mice were placed in the home arm and allowed to explore the Y-maze freely and without any optogenetic manipulation during habituation for 5 minutes. This was followed by 5 days of training, where mice can still move freely through the maze for 5 minutes, but entering the paired arm triggered optogenetic inhibition (constant light, 6mW) of AgRP neurons. On the test day, mice received no stimulation when entering the paired arm and movement was recorded for 5 minutes and analysed with Ethovision 15XT (Noldus). Y-maze was cleaned and wiped with 30% ethanol between animals.

Immunohistochemistry: After completion of experiments mice were perfused and brains stored in 4%PFA for 48 hours before transferred into 30% sucrose. Brain sections at 40 microns were collected in sets of 4 using a cryostat and 1 set used for immuno histological verification of viral injection (chicken anti GFP 1:1000 (ab13970, Abcam), Alexa anti chicken 465nm (Invitrogen) and fiber placement as described before^17^.

Statistical Analysis: Graphs were generated and statistical analysis performed using GraphPad Prism for MacOS X. Data are represented as mean ± SEM. Two-way ANOVAs with post hoc tests were used to determine statistical significance (< 0.05 was considered statistically significant and is indicated on figures and in figure legends). Post hoc analysis of fiber photometry peri-events was performed using bootstrapped confidence intervals 64with following parameters: 95 % confidence intervals, 1000 resampling, 1000 permutations and 0.5 seconds consecutive threshold.

## Acknowledgement

This research was funded by the Australian Government through the Australian Research Council (ARC). Alex Reichenbach is the recipient of an ARC Australian Discovery Early Career Award (DE240100950) funded by the Australian Government. Components of figures were created with BioRender.com

---

**Extended Data Figure 1.**
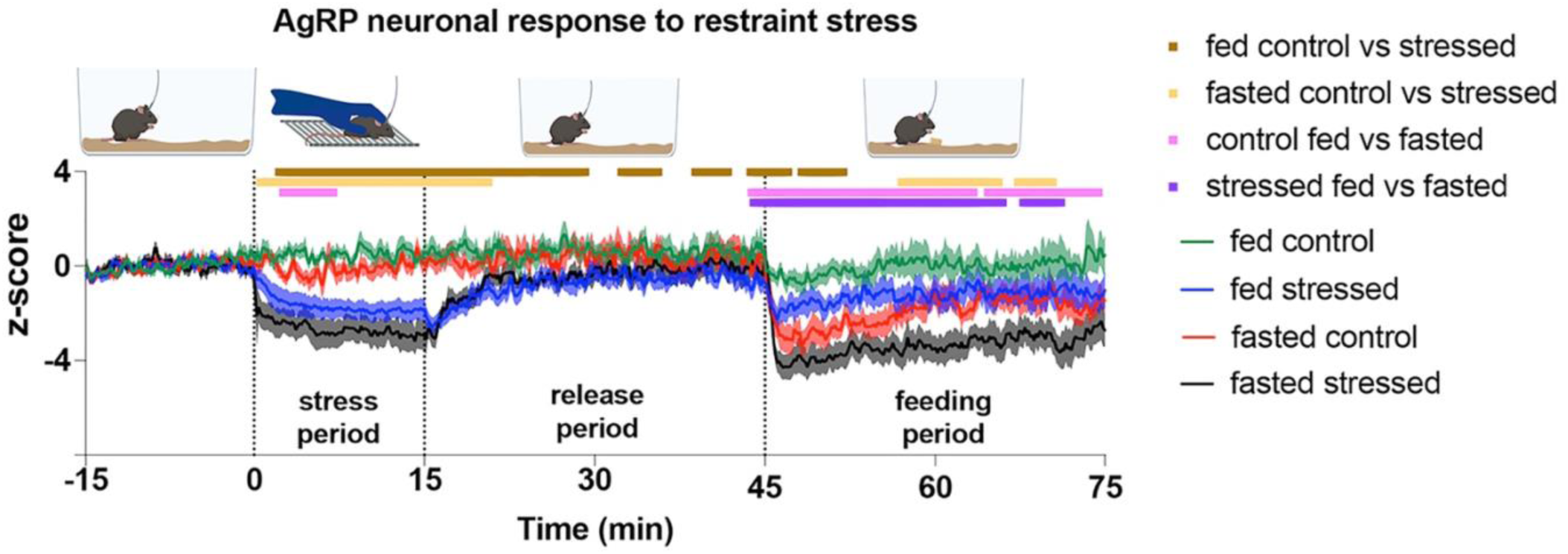
Between group comparisons of data shown in Fig1 J: AgRP neuronal activity over 90-minute ecording showing response to 15-minute restraint stress and 30-minute food access relative to a pre-stress baseline period. N= 11 fed control; N= 9 fasted control; N= 12 fed stressed; N= 10 asted stressed. Significant differences are indicated by the lines above graphs as calculated by bootstrapped confidence intervals with 1000 resamples at 95%confidence and 0.5sec consecutive threshold

**Extended Data Figure 2.**
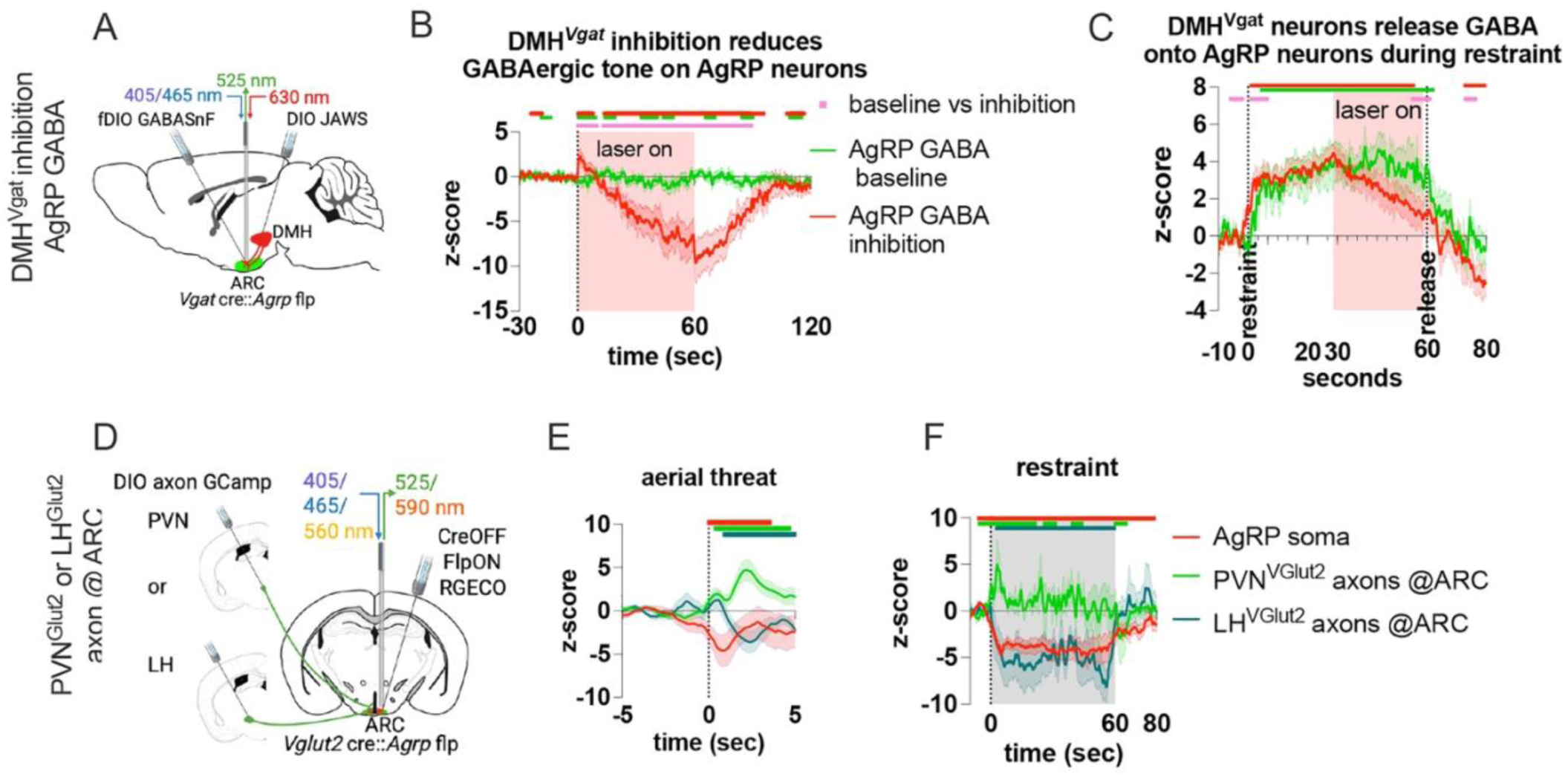
A) Schematic of the viral approach to inhibit DMH*Vgat* neurons while measuring GABA release onto AgRP neurons. B) GABA release onto AgRP neurons during rest in home cage with (red) and without (green) DMH*Vgat* neuron inhibition C) GABA release onto AgRP neurons during 1 minute restraint stress with 30 second DMH*Vgat* neuronal inhibition during the second half of restraint. D) viral approach to target glutamatergic input onto AgRP neurons from PVN and LH Vglut2 neurons. PVN*Vglut2* ARC axonal (green) and PVN*Vglut2* ARC axonal (cyan) and AgRP neuronal (red) response to aerial threat (E) and to one minute restraint (F). N (Vgat JAWS) =3, N(Vglut2 axons) = 2 each. Significant differences are indicated by the lines above graphs as calculated by bootstrapped confidence intervals with 1000 resamples at 95%confidence and 0.5sec consecutive threshold, with the same colours indicating differences from baseline. Pink lines in B and C indicated differences between groups.

**Extended Data Figure 3.**
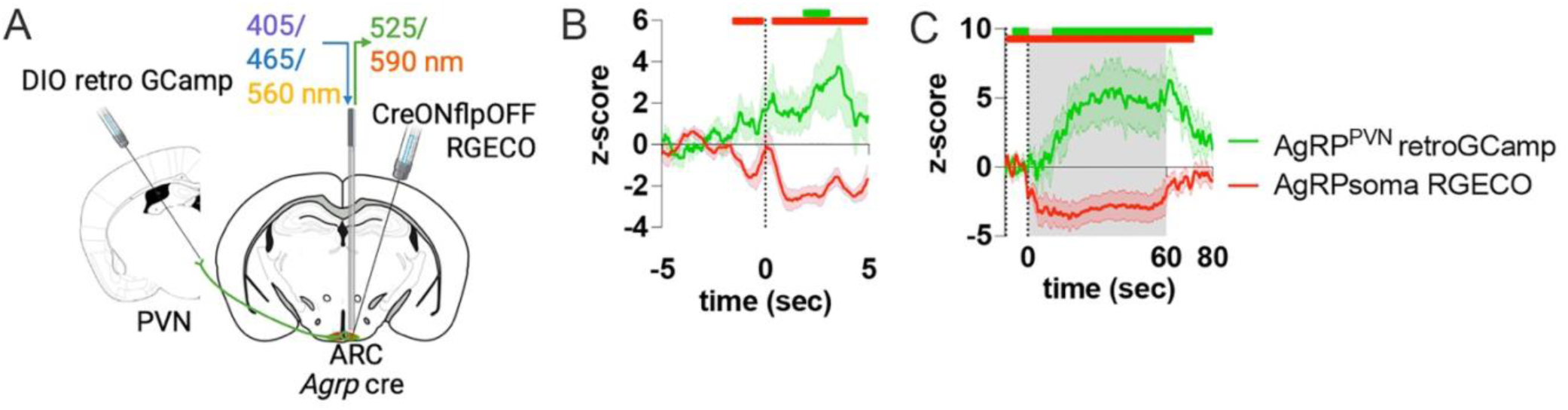
A) Schematic of viral approach to simultaneous record cell bodies from AgRP->PVN projecting neurons and average AgRP soma activity in response to aerial threat (B) and one minute restraint C). N = 3. Significant differences are indicated by the lines above graphs as calculated by bootstrapped confidence intervals with 1000 resamples at 95%confidence and 0.5sec consecutive threshold, with the same colours indicating differences from baseline.

| Figure 1 | Statistical test | Group n | Main analysis result | Significant post-hoc test |
| --- | --- | --- | --- | --- |
| 1C | 2way ANOVA | N= 6 | Metabolic state p=0.4528<br>Transition p =0. 8367 |  |
| 1D | Bootstrapped CI intervals + permutation test |  |  |  |
| 1E | Bootstrapped CI intervals + permutation test |  |  |  |
| 1F | Bootstrapped CI intervals + permutation test |  |  |  |
| 1G | Bootstrapped CI intervals + permutation test |  |  |  |
| 1I | Bootstrapped CI intervals + permutation test | N= 10 fed<br>N=6 fasted |  |  |
| 1J/ Supp Fig1 | Bootstrapped CI intervals + permutation test | N= 11 fed control<br>N= 9 fasted control<br>N= 12 fed stressed<br>N= 10 fasted stressed | <u>Fed -stressed vs control:</u><br>Baseline p = 0.241<br>Comparison p =0.006<br><u>Fasted -stressed vs control:</u><br>Baseline p = 0.453<br>Comparison p =0.008<br><u>control -fed vs fasted:</u><br>Baseline p = 0.631<br>Comparison p =0.037<br><u>stressed -fed vs fasted:</u><br>Baseline p = 0.842<br>Comparison p =0.076 |  |
| 1K | 2way ANOVA |  | Metabolic state p<0.0001<br>treatment p =0. 7442 | Uncorrected Fisher's LSD<br>Fed- control vs stressed p =0.6640<br>Fasted – control vs stressed p = 0.9835<br>**Control – fed vs fasted p<0.0001**<br>**Stressed – fed vs fasted p = 0.0001** |
| 1L | Bootstrapped CI intervals + permutation test |  | Baseline p = 0.616<br>Comparison p =0.346 |  |
| 1M | Bootstrapped CI intervals + permutation test |  | <u>Fed -control vs stressed:</u><br>Baseline p = 0.57<br>Comparison p =0.001<br><u>Fasted -control vs stressed:</u><br>Baseline p = 0.171<br>Comparison p =0.003 |  |

| Figure 2 | Statistical test | Group n | Main analysis result | Significant post-hoc test |
| --- | --- | --- | --- | --- |
| 2B | Bootstrapped CI intervals + permutation test | N=7 |  |  |
| 2C | Bootstrapped CI intervals + permutation test |  |  |  |
| 2D | Bootstrapped CI intervals + permutation test |  |  |  |
| 2F | Bootstrapped CI intervals + permutation test | N=3 |  |  |
| 2G | Bootstrapped CI intervals + permutation test |  |  |  |
| 2H | Bootstrapped CI intervals + permutation test |  |  |  |
| 2J | Bootstrapped CI intervals + permutation test | N=3 |  |  |
| 2K | Bootstrapped CI intervals + permutation test |  |  |  |
| 2L | Bootstrapped CI intervals + permutation test |  |  |  |
| 2N | Bootstrapped CI intervals + permutation test | N=5 | Baseline p = 0.306<br>Comparison p <0.001 |  |
| 2O | Bootstrapped CI intervals + permutation test |  |  |  |
| 2Q | Bootstrapped CI intervals + permutation test | N=3 |  |  |
| 2P | Bootstrapped CI intervals + permutation test |  |  |  |

| Figure 3 | Statistical test | Group n | Main analysis result | Significant post-hoc test |
| --- | --- | --- | --- | --- |
| 3D | Bootstrapped CI intervals + permutation test | N=4 |  |  |
| 3E | Bootstrapped CI intervals + permutation test |  |  |  |
| 3F | Bootstrapped CI intervals + permutation test |  |  |  |
| 3H | Bootstrapped CI intervals + permutation test | N=4 |  |  |
| 3I | Bootstrapped CI intervals + permutation test |  |  |  |
| 3J | Bootstrapped CI intervals + permutation test |  |  |  |
| 3L | Bootstrapped CI intervals + permutation test | N=4 |  |  |
| 3M | Bootstrapped CI intervals + permutation test |  |  |  |
| 3N | Bootstrapped CI intervals + permutation test |  |  |  |
| 3P | Bootstrapped CI intervals + permutation test | N=4 |  |  |
| 3Q | Bootstrapped CI intervals + permutation test |  |  |  |
| 3R | Bootstrapped CI intervals + permutation test |  |  |  |
| 3T | Bootstrapped CI intervals + permutation test | N=4 |  |  |
| 3U | Bootstrapped CI intervals + permutation test |  |  |  |
| 3V | Bootstrapped CI intervals + permutation test |  |  |  |
| 3X | Bootstrapped CI intervals + permutation test | N=3 |  |  |
| 3Y | Bootstrapped CI intervals + permutation test |  |  |  |

| Figure 4 | Statistical test | Group n | Main analysis result | Significant post-hoc test |
| --- | --- | --- | --- | --- |
| 4C | 2way ANOVA | N=7 control<br>N=14 JAWS | side p=0.4395<br>**genotype p =0. 0467** | Uncorrected Fisher's LSD<br>Control arm- WT vs JAWS p =0.0859<br>Paired arm- WT vs JAWS p = 0.2594<br>WT – control vs paired p = 0.8343 |
|  |  |  |  | JAWS – control vs paired p=0.2980 |
| **4D** | 2way ANOVA |  | side p=0.7788<br>**genotype p =0. 0074** | Uncorrected Fisher's LSD<br>**Control arm- WT vs JAWS p =0.0132**<br>Paired arm- WT vs JAWS p = 0.1696<br>WT – control vs paired p = 0.4926<br>JAWS – control vs paired p=0.6270 |
| **4E** | 2way ANOVA |  | **side p=0.0462**<br>genotype p =0. 3373 | Uncorrected Fisher's LSD<br>**Control arm- WT vs JAWS p =0.0415**<br>Paired arm- WT vs JAWS p = 0.4681<br>WT – control vs paired p = 0.9748<br>**JAWS – control vs paired p=0.0017** |
| **4F** | 2way ANOVA |  | side p=0.0556<br>genotype p =0. 2199 | Uncorrected Fisher's LSD<br>Control arm- WT vs JAWS p =0.9416<br>Paired arm- WT vs JAWS p = 0.0874<br>WT – control vs paired p = 0.6400<br>**JAWS – control vs paired p=0.0036** |
| **4G** | 2way ANOVA |  | **side p=0.0172**<br>genotype p =0. 2601 | Uncorrected Fisher's LSD<br>Control arm- WT vs JAWS p =0.0724<br>Paired arm- WT vs JAWS p = 0.8185<br>WT – control vs paired p = 0.5350<br>**JAWS – control vs paired p=0.0015** |
| **4H** | 2way ANOVA |  | **side p=0.0073**<br>genotype p =0. 5128 | Uncorrected Fisher's LSD<br>Control arm- WT vs JAWS p =0.1937<br>Paired arm- WT vs JAWS p = 0.6996<br>WT – control vs paired p = 0.3265<br>**JAWS – control vs paired p=0.0012** |

| **Suppl Figure 2** | **Statistical test** | **Group n** | **Main analysis result** | **Significant post-hoc test** |
| --- | --- | --- | --- | --- |
| **2B** | Bootstrapped CI intervals + permutation test | N=3 | Baseline p = 0.493<br>Comparison p =0.008 |  |
| **2C** | Bootstrapped CI intervals + permutation test |  | Baseline p = 0.864<br>Comparison p =0.486 |  |
| **2E** | Bootstrapped CI intervals + permutation test | N=2 PVN<br>N=2 LH |  |  |
| **2F** | Bootstrapped CI intervals + permutation test |  |  |  |

| **Suppl Figure 3** | **Statistical test** | **Group n** | **Main analysis result** | **Significant post-hoc test** |
| --- | --- | --- | --- | --- |
| **3B** | Bootstrapped CI intervals + permutation test | N=3 |  |  |
| **3C** | Bootstrapped CI intervals + permutation test |  |  |  |

## Notes

### Competing Interest Statement

The authors have declared no competing interest.

